## Supporting Information for "PROTAC-induced Protein Structural Dynamics in Targeted Protein Degradation"

### Supporting Method

#### Modeling CRBN-dBETx-BRD4$^{BD1}$ ternary complexes

The inhibitor (JQ1)-bound BRD4$^{BD1}$ crystal structure (PDB ID: 3MXF)[1] and pomalidomide-bound CRBN crystal structure (PDB ID: 4CI3)[2] were downloaded from the PDB website. Four BRD4$^{BD1}$ PROTACs (dBET1, dBET23, dBET57 and dBET70) were prepared using the builder program in the Molecular Operating Environment (MOE). Ternary complex ensembles were generated using the method 4B protocol implemented in the MOE program (Chemical Computing Group)[3]. CRBN-dBETx-BRD4$^{BD1}$ ternary complex was modelled in four steps: (1) CRBN-BRD4$^{BD1}$ protein-protein docking with a conventional global protein-protein docking approach; (2) Robust conformational sampling of isolated dBET; (3) CRBN-BRD4$^{BD1}$ pre-generated poses and dBET conformation alignment with respective inhibitors in either protein kept intact while using Maximum Common Substructure (MCS) approach to determine the match on-the-fly between the binding ligands in the protein-protein docking ensemble and supplied dBET; (4) scoring and clustering of modelled CRBN-dBETx-BRD4$^{BD1}$ ternary complex. Before performing protein–protein docking in step 1, two crystal structures were prepared by adding missing hydrogens and assigning an appropriate protonation state for each protein. In the protein–protein docking step 1, two binary complexes interacted without a linker connecting two warheads. The protein–protein poses were generated, and conformations were stored for the next step. In the dBET (or PROTAC in general) conformational sampling step 2, five dBETs were provided for robust conformational searching. Each iteration was set to 10000. Ligand core root mean square deviation (RMSD) was set as the default. In the dBETx conformational searching process, each binding warhead in the dBETs was held rigid to retain its bound conformation, whereas the linker region was simulated on-the-fly. A MCS approach was applied in the program to maximize dBET warheads alignment with the binding warheads. The simulated dBET conformations were then stored for the next step. In the protein–protein pose and dBET conformation alignment step, protein–protein poses with two warheads in each binding pocket were aligned with the dBETs' two binding moieties. The generated ternary complex CRBN-dBETx-BRD4$^{BD1}$ ensembles were minimized, scored, clustered. Selected represented conformations with docking scores were stored in pdb format for further visualization analysis.

**Constructing CRL4A E3 ligase scaffold: DDB1/CUL4A/NEDD8/Rbx1/E2/Ub/CRBN**

We obtained 12 DNA damage-binding protein 1 (DDB1) PDB structures from the PDB website and named them by the index ID: A1_DDB1 (PDB ID: 4E54), A2_DDB1 (PDB ID: 3E0C), A3_DDB1 (PDB ID: 3I8E), A4_DDB1 (PDB ID: 4TZ4), A5_DDB1 (PDB ID: 4A0L), B1_DDB1 (PDB ID: 2B5L), B2_DDB1 (PDB ID: 3EI4), B3_DDB1 (PDB ID: 3EI3), B4_DDB1 (PDB ID: 6PAI), C1_DDB1 (PDB ID: 6FCV), C2_DDB1 (PDB ID: 4A08), and C3_DDB1 (PDB ID: 4A0B). CUL4A with its DDB1 crystal structure was downloaded (PDB ID: 2HYE) and used as a template. 12 DDB1 crystal structures were superimposed on the CUL4A/DDB1 template via β-propeller domains (BPB) to build 12 DDB1/CUL4A structures. CRBN/DDB1 from A4_DDB1 (PDB ID: 4TZ4) was used as a template, and the rest of the DDB1/CUL4A structures were superimposed via the BPA/BPC domain to build 12 DDB1/CUL4A/CRBN complexes. Next, E2D1 (PDB ID 5FER chain B) was aligned to (PDB ID 6TTU) to replace E2D2. We then aligned the 12 CRBN/DDB1/CUL4A complexes via residues in the C-terminal domain (residues from 416 to 672) in the CUL4A. Components having NEDD8, Rbx1, E2 and Ub with CUL1 (PDB ID 6TTU) were added to the above 12 CRBN/DDB1/CUL4A structures. The final 12 DDB1/CUL4A/Rbx1/NEDD8/E2/Ub/CRBN E3 ligase scaffolds were ready for the next step (**Figure S3 for flowchart**). Next, we analyzed and classified the 12 CRL4A E3 ligase scaffolds DDB1/CUL4A/NEDD8/Rbx1/E2/Ub/CRBN based on the interface distance between the CRBN E3 ligase and the E2. Specifically, we measured residue distances from CRBN E3 to the E2 interface plane among these 12 CRL4A E3 ligase scaffolds. If the CRBN E3 ligase residue distances to E2 plane were < 10 Å, with no overlapping or clashing, we grouped those CRL4A E3 ligase scaffolds as cluster A; if the CRBN E3 ligase residue distances to E2 plane were > 10 Å (the exact gap distance was from ~14 Å to ~34 Å), we grouped those scaffolds as cluster B. If CRBN E3 and E2 overlapped, we grouped those scaffolds as cluster C, considering them as clashing and abandoned them (**Figure S4)**.

**Degradation machinery complex assembling: DDB1/CUL4A/NEDD8/Rbx1/E2/Ub/CRBN-dBETx-BRD4$^{BD1}$**

CRBN-dBETx-BRD4$^{BD1}$ conformations generated from protein–protein docking were aligned with the CRL4A E3 ligase scaffolds using clusters A and B (total of nine CRL4A E3 ligase scaffolds) via the common CRBN structure. Because there are 12 lysine residues on the surface of

BRD4$^{BD1}$ (**Figure S1A**), we considered all the lysine residues that potentially lead to ubiquitination by measuring the distance between the oxygen from the C-terminal glycine of the ubiquitin and the nitrogen atom of surface-exposed lysine residue(s) of the BRD4$^{BD1}$ by using VMD[31] (**Table S1**). Among the modelled CRBN-dBETx-BRD4$^{BD1}$ conformations each with CRL4A E3 ligase scaffolds (A1, A2, A3, A4, A5, B1, B2, B3, B4), only those with the measured distance less than or close to 16 Å without clashing were kept for further study, as suggested by previous study[25].

**MD simulations**

The top selected CRBN-dBETx-BRD4$^{BD1}$ conformations from combining protein–protein docking, and structural based steps were used for MD simulations to capture the dynamics of CRBN-dBETx-BRD4$^{BD1}$ and degradation machinery complexes (**Table S1**). For each dBET, we selected one CRBN-dBETx-BRD4$^{BD1}$ conformation. For reference, we also investigated and performed MD simulations for resolved crystal structures. Among four studied dBETs in this work, only CRBN-dBET23-BRD4$^{BD1}$ (PDB ID 6BN7) has PDB structures for starting the MD simulations. Other ternary complexes missing the CRBN E3 component (PDB ID 4ZC9, no dBET1[4]) or the dBET degrader (PDB ID 6BNB, no dBET57, PDB ID 6BN9 no dBET70) were not considered. Before proceeding to MD simulations, the missing part in each protein component of each CRL4A E3 ligase complex was built by using structural alignment or the homology modeling method[5]. In brief, we used (PDB ID 4A0L) as a template for A1_DDB1, (PDB ID 3L7K) as a template for A3_DDB1, and (PDB ID 5HTB) as a template for A5_DDB1 in homology modeling. For the remaining CRL4A E3 ligase complexes (A2, A4, B1, B2, B3, B4), we built the missing part directly using the homology modeling method. We chose one CRL4A E3 ligase complex from each category: A1 in category A and B1 in category B based on the high populations in that complex. Then two criteria were considered to select the top CRBN-dBETx-BRD4$^{BD1}$ for the ternary complex MD simulations and for degradation machinery complex MD simulations: 1) the common CRBN-dBETx-BRD4$^{BD1}$ shared in the category (**Figure S1 and Figure S5**) the best protein–protein docking score (**Table S2**).

MD simulations were performed using the AMBER20 software package[6] with AMBER ff14SB force field[7] for proteins and GAFF2[8] for ligand parameterization. The atomic charges were computed using AM1-BCC[9]. A standard AMBER simulation protocol was used. The sequential minimization steps were performed for the hydrogen atoms, side chains and the entire protein complex to clean any clashes. The system was solvated in a TIP3P water box[10] with distance 12 Å from the edge of the protein. To neutralize the CRL4A E3 ligase complex system, 40 and 46 $Na^+$ ions were added to the water box for C4 cluster and for C12, respectively, with no counter ion added to the ternary complex. Subsequently, the whole complex system with water was minimized by only focusing on water molecules, then the entire complex system. Heating of the complex system from 50 K to 300 K with increments of 25 K was achieved in 200-ps NPT simulation (time step of 2 fs) for each temperature. The system was subsequently equilibrated at 300 K for 500-ps NPT simulation with all atoms relaxed. Long-range electrostatic interactions were computed by the particle mesh Ewald method (PME)[11]. The SHAKE constraint was applied to all atoms including hydrogens.[12] The Langevin thermostat with a damping constant of 2 $ps^{-1}$ was used to maintain a temperature of 300 K. The NPT ensemble at 300 K was used for MD production run for 400 ns, and each frame was collected at 1-ps time intervals. We generated eight CRBN-dBETx-$BRD4^{BD1}$ complex structures and performed 3 MD runs with different initial velocities for each structure. Sixteen degradation machinery complex structures were constructed, and we ran one MD for each complex (**Table 2**).

**Dihedral angle correlation**

We used the T-analyst program[13] to calculate the pairwise correlations between the pseudo dihedral angles and the PROTAC dihedral angles. Each side-chain dihedral angle was recorded every 50 ps through 400 ns trajectories to generate 8000 different angles per dihedral selection. Pairwise correlations were computed using a Pearson correlation formula. We converted the side-chain dihedral angles to Cartesian coordinates by means of equations (1)-(3) to accurately capture their differences and means, thereby preventing erroneous computation of their correlation at the discontinuity margin (±180° or 360°/0°)[13,14]. Notably, a positive correlation between two sidechains indicates that the two sides rotate similarly during MD simulation.

$$r_{xy} = \frac{\sum_{i=1}^{n}(x_i-\bar{x})(y_i-\bar{y})}{\sqrt{\sum_{i=1}^{n}(x_i-\bar{x})^2}\sqrt{\sum_{i=1}^{n}(y_i-\bar{y})^2}} \quad (1)$$

$$\bar{x} = \arctan\left(\frac{sin(x_1)+sin(x_2)+\cdots+sin(x_n)}{cos(x_1)+\cos(x_2)+\cdots+cos(x_n)}\right) \quad (2)$$

$$x_i - \bar{x} = \arctan\left(\frac{\sin(x_i)\cos(\bar{x})-\sin(\bar{x})\cos(x_i)}{\cos(x_i)\cos(\bar{x})+\sin(x_i)\sin(\bar{x})}\right) \quad (3)$$

$r_{xy} = \ Dihedral\ Pearson\ Correlation, \overline{x} = mean\ of\ dihedral\ angles,$

$x_i = side\ chain\ dihedral\ angles$

**Interaction energy calculations**

We used MM/PBSA method from AMBER20[15] to evaluate the stability of ternary complex by computing the interaction energy between protein (CRBN-BRD4$^{BD1}$) and ligand (dBETs). From a total of 8,000 MD frames making up the 400-ns ternary complex trajectories, system conformations were analyzed every 4 ns. This method computes the energy (E) of a system from the protein (CRBN-BRD4$^{BD1}$), ligand (dBETs) and protein–ligand complex (CRBN–dBETs–BRD4$^{BD1}$), and computes the interaction energy as:

$$\Delta E = \ E_{\mathrm{CRBN-dBETs-BRD4}^{BD1}} - E_{\mathrm{CRBN-BRD4}^{BD1}} - E_{\mathrm{dBETs}} \ (4)$$

The solute dielectric is set to 15.0 to consider the polar surface of protein and solvent dielectric was set to 80.0.

**Pairwise interaction network**

Gromacs force distribution analysis (FDA) was used to compute nonbonding forces with Leonard-Jones' potential and Coulomb potential[16].The parameter 10 Å was used for the short-range interaction cutoff. The long-range electrostatic forces were computed with PME[11].The first 40 ns of MD simulation were treated as equilibrium plus, and thus the FDA was performed with the following 360 ns. We calculated the sum of pair-wise forces of the degradation complex by using our in-house script. In addition, we constructed the pair-wise forces network of the intramolecular attraction of degradation complex to visualize the shortest path of the PROTAC-guided essential motion at the hinge region by using python library Networkx[17] (**Figure S14**). To reduce the complexity of the network, we eliminated pair-wise forces that had minor attraction or repulsion. Specifically, pair-wise forces that were within -10 and 10 pN were eliminated.

**Principle component analysis (PCA)**

To observe major protein motions, we performed PCA using CPPTRAJ and our in-house code[18,19]. PCA of MD trajectories for all the simulated systems involved using backbone atoms in the full degradation machinery complex. The first and second PCs were analyzed to reveal the dominant motions (**Figure S13**).

### Data Availability

The modeled structures of the four degradation complexes are available in supplementary files: dbet1_initial.pdb, dbet23_intial.pdb, dbet57_initial.pdb, and dbet70_initial.pdb. Other files reported in this study, including the input, output, parameter, and trajectories files, are available upon request.

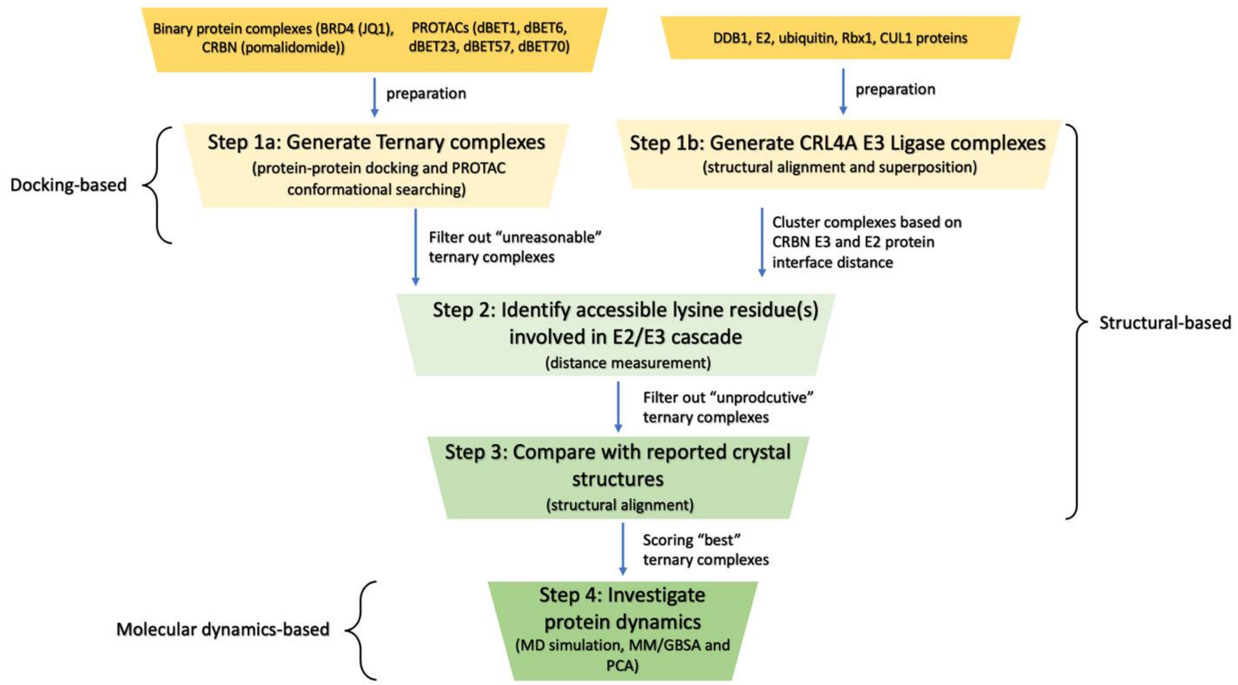


**Scheme S1:** The overall workflow of this study includes the docking-based, structural-based, and molecular dynamics (MD)-based approaches.

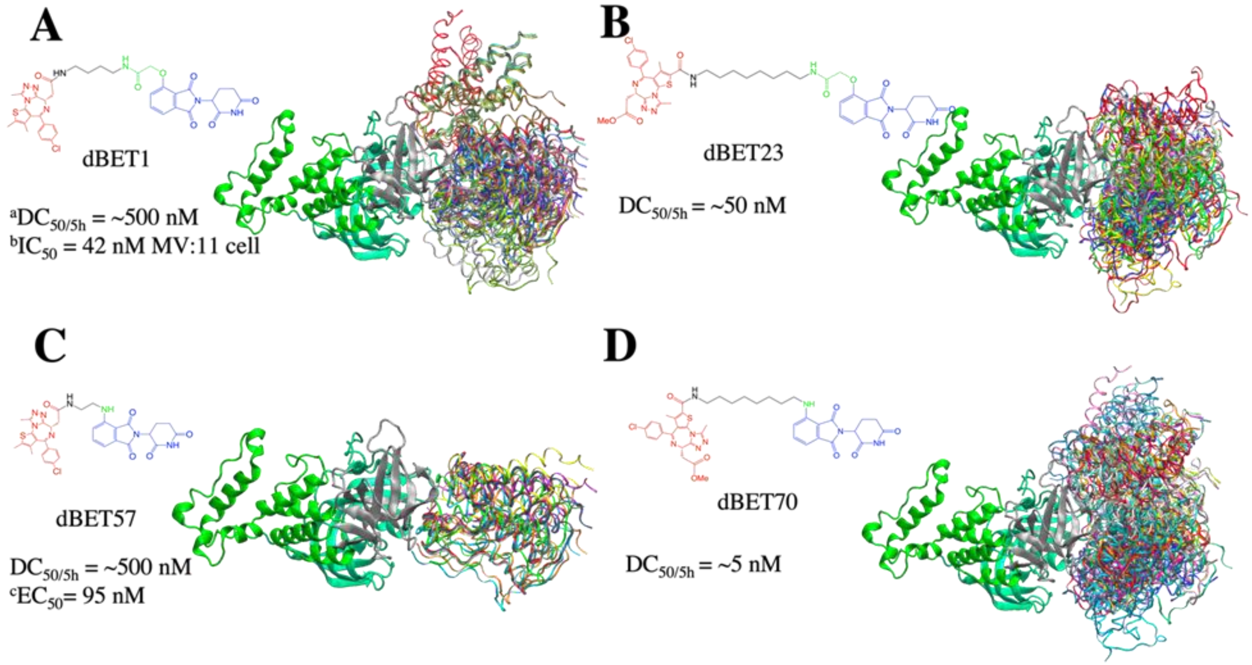


**Figure S1: Conformational ensembles of ternary complexes (CRBN-degrader of bromodomain and extra-terminal domain (dBETx)-$BRD4^{BD1}$) from protein–protein docking.** All protein–protein docking conformations for each PROTAC were superimposed based on CRBN (thick, ribbon) to show different poses of $BRD4^{BD1}$ (rainbow color; ribbon). Each dBET PROTAC has multiple binding poses that contribute to different degradation efficiency. **(A)** dBET1 with 186 ternary complex conformations. **(B)** dBET23 with 168 ternary complexes. **(C)** dBET57 with 24 ternary complexes. **(D)** dBET70 with 183 ternary complexes. [a]Degradation profile $DC_{50/5h}$ for four PROTACs was obtained from EGFP/mCherry reporter assay published in reference #21. [b]Data published in reference #20[20]. [c]Data published in reference #21.

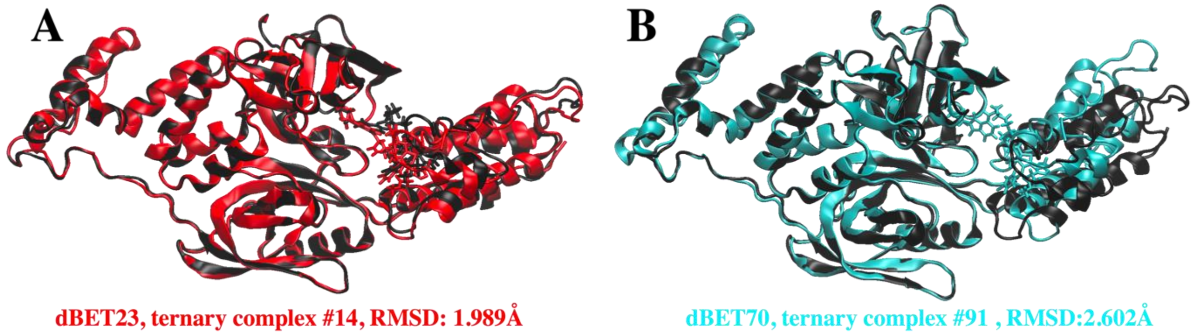


**Figure S2**. **Superpositions of predicted CRBN-dBETx-BRD4$^{BD1}$ ternary complexes (red for dBET23 and cyan for dBET70) with reported crystal structures (Black). (A)** Predicted CRBN-dBET23-BRD4$^{BD1}$ ternary complex index #14 (red) has RMSD of 1.99 Å with respect to the crystal structure (PDB ID: 6BN7). **(B)** Predicted CRBN-dBET70-BRD4$^{BD1}$ ternary complex index #91 (cyan) has RMSD of 2.60 Å with respect to the crystal structure (PDB ID: 6BN9). Note, dBET70 degrader was not resolved in the crystal structure. The structural superposition is performed based on the backbone of CRBN and RMSD calculation is conducted based on the backbone of BRD4$^{BD1}$. PROTAC dBET23 and dBET70 are shown in licorice representation.

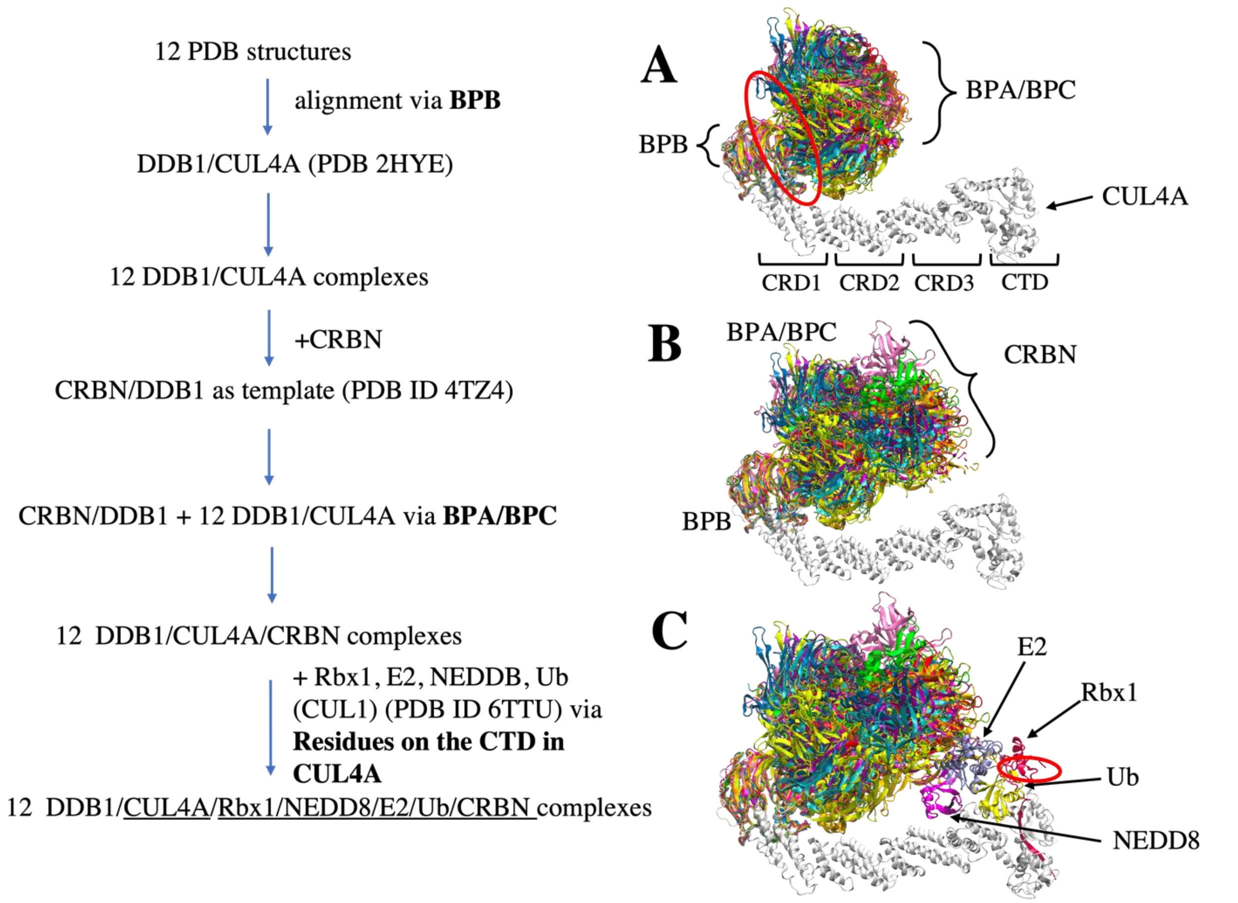


**Figure S3: Flowchart of CRL4A E3 ligase scaffolds construction**. **(A)** structure alignments of 12 DDB1 PDB structures to the CUL4A based on the BPB part. Red circle indicates a hinge loop connecting BPB and BPA/BPC in DDB1. **(B)** Structure alignment of CRBN E3 ligase to the DDB1/CUL4A based on the BPA/BPC part, resulting 12 DDB1/CUL4A/CRBN scaffolds. **(C)** E2/Rbx1/Ub/NEDD8 components (PDBID: 6TTU) are added to the /DDB1/CUL4A/CRBN scaffolds to make 12 diverse CRL4A E3 ligase scaffolds. Red circle indicates another hinge loop in the Rbx1.

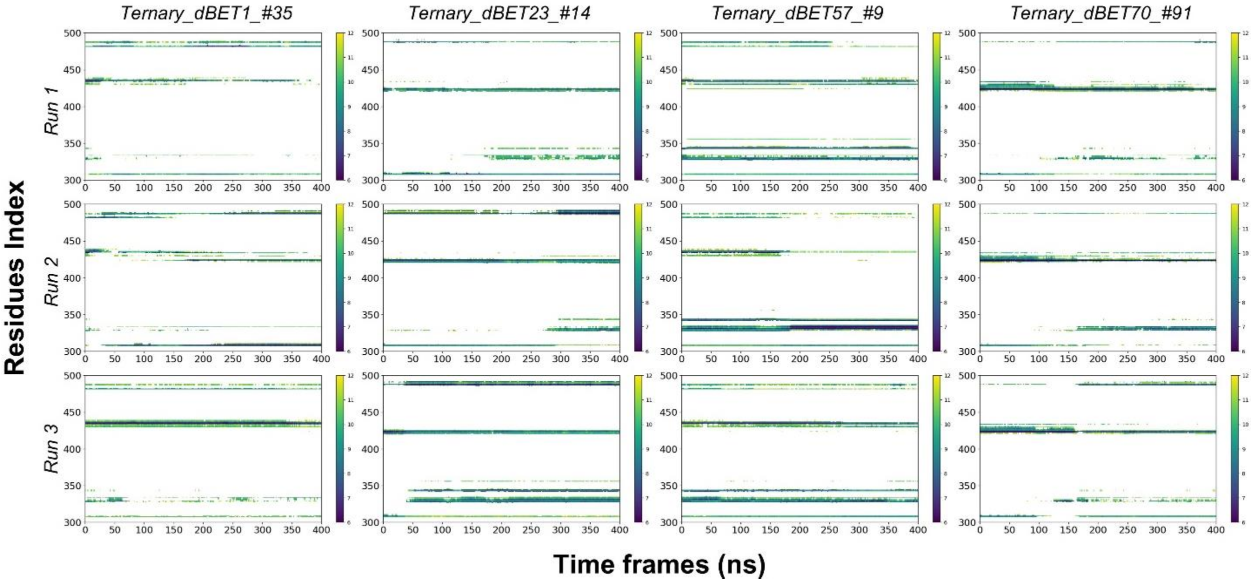


**Figure S4. Residues Contact map between PROTACs and BRD4$^{BD1}$ throughout 400ns of MD simulation.** X-axis is the residues index of BRD4$^{BD1}$ and y-axis is the Time of MD simulation. The distance cutoff is 12Å.

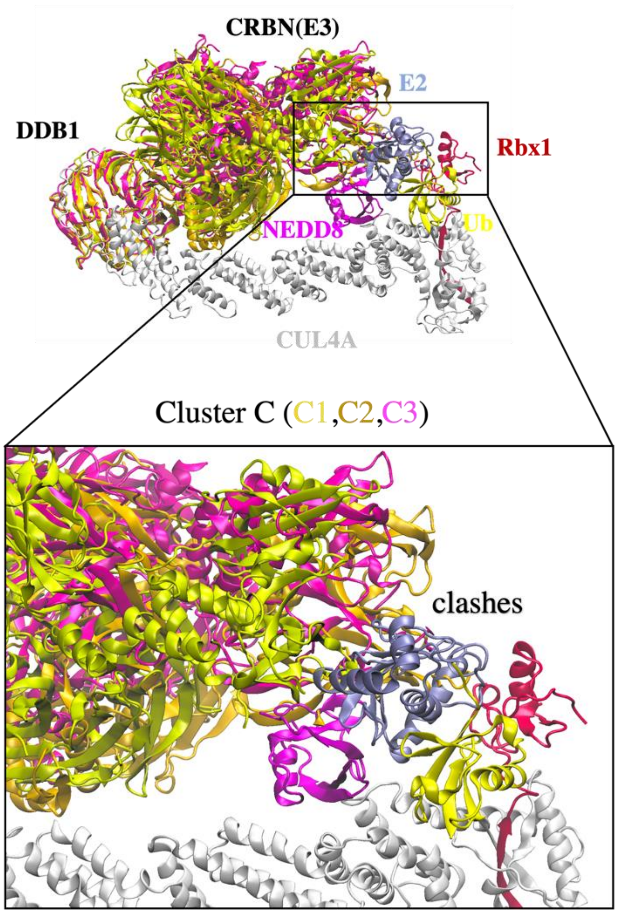


**Figure S5. Cluster C with clashes between CRBN E3 and E2.** Three DDB1 conformations (C1, yellow; C2, gold; and C3, pink) were found to have protein clashing and is not considered for further degradation machinery complex construction.

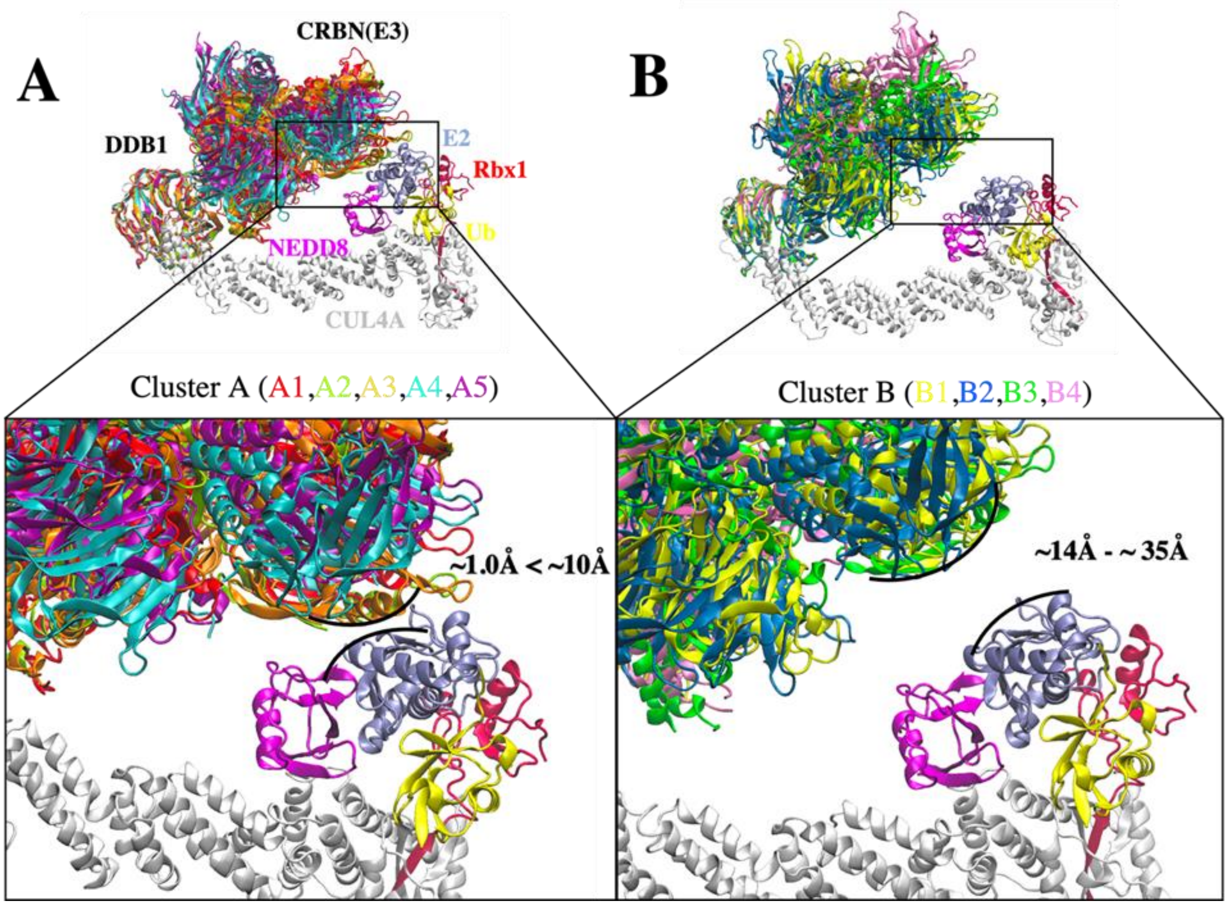


**Figure S6: Conformational clusters of CRL4A E3 ligase scaffolds. (A)** Cluster A with interface distance between E2 and E3 ligase ranges from ~1.0 Å to ~10 Å. Five DDB1 conformations were analyzed and grouped in cluster A. A1 (red), A2 (mint), A3 (orange), A4 (cyan), and A5 (purple). **(B)** Cluster B with interface distance between E2 and E3 ligase ranges from ~14 Å to ~35 Å. Four DDB1 conformations were analyzed and grouped in cluster B. B1 (yellow), B2 (blue), B3 (green), and B4 (pink). Black curved lines indicate the E2–E3 interface distance in each cluster with the closest distance. All nine DDB1 conformations were aligned based on β-propeller domains in DDB1. Other protein components: CUL4A (white), NEDD8 (magenta), E2 (ice blue), Ub (yellow), and Rbx1 (red) were taken from PDB ID 6TTU and added to nine clusters to construct nine CRL4A E3 ligase scaffolds.

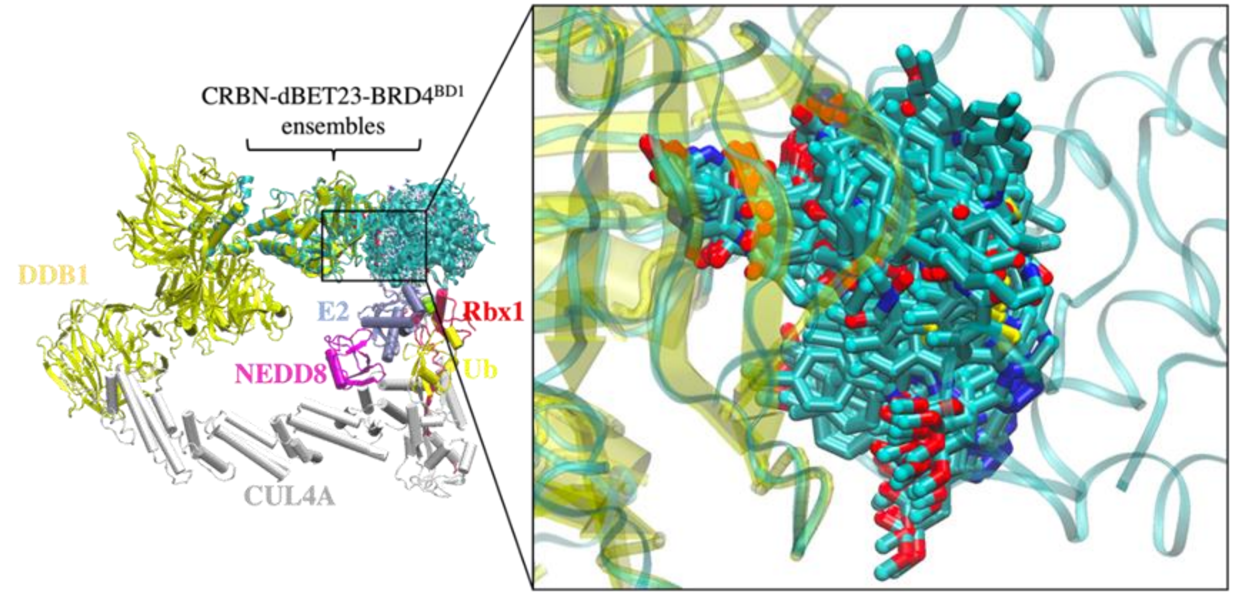


**Figure S7: Conformational ensemble of dBET23 from protein–protein docking.** 168 CRBN-dBET23-BRD4$^{BD1}$ ternary complexes were applied to cluster B1 to construct 104 degradation machinery complexes. Ternary complexes were superimposed on the CRL4A E3 ligase by aligning the backbone of CRBN. The zoom-in region shows the flexibility of the linker of dBET23.

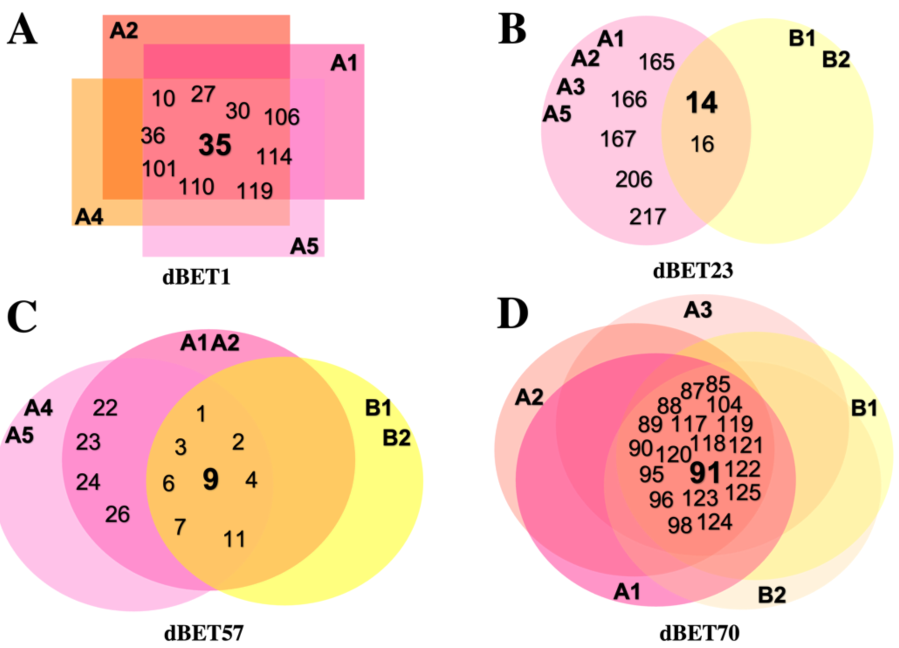


**Figure S8: Overlapping diagram representation of common CRBN-dBETx-BRD4$^{BD1}$ ensembles found in each degradation machinery complex**. Numbers represent ensemble index. **(A)** CRBN-dBET1-BRD4$^{BD1}$ ensembles fall within A1, A2, A4 and A5 clusters. **(B)** CRBN-dBET23-BRD4$^{BD1}$ ensembles fall within A1, A2, A3, A5, B1, and B2 clusters. **(C)** CRBN-dBET57-BRD4$^{BD1}$ ensembles fall within A1, A2, A4, A5, B1, and B2 clusters. **(D)** CRBN-dBET70-BRD4$^{BD1}$ ensembles fall within A1, A2, A3, B1, and B2 clusters.

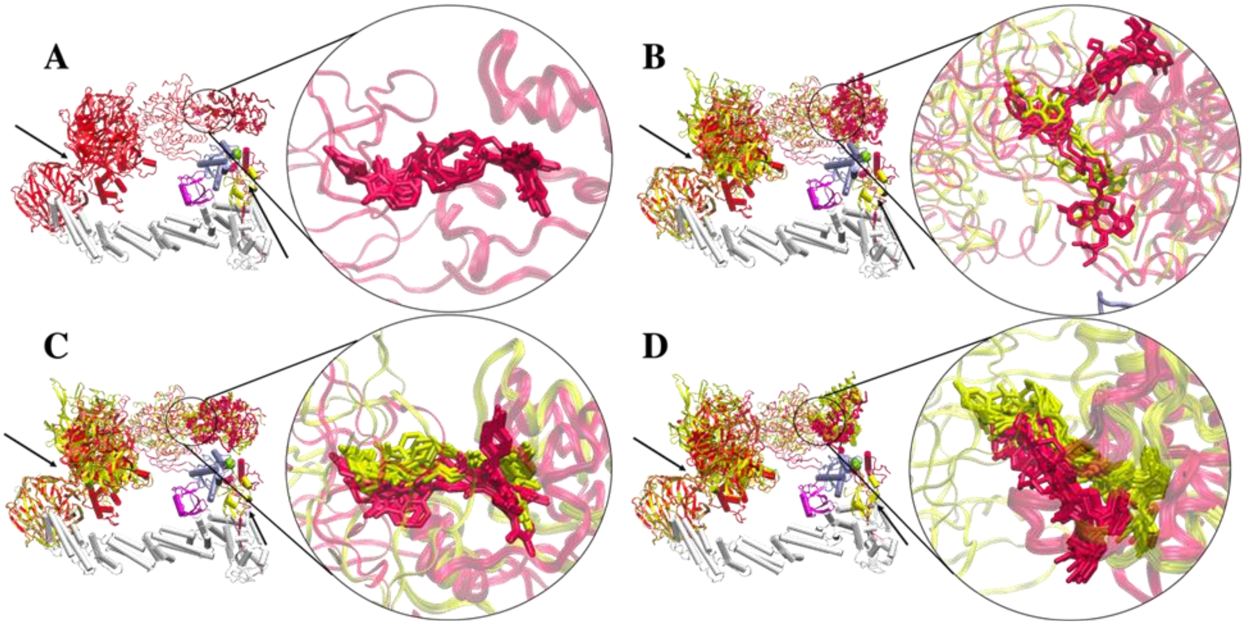


**Figure S9. Clusters of PROTAC conformations from top-ranked docking scores.** Aligning CRBN of the two representative clusters A1 and B1 shows similar PROTAC conformations. **(A)** dBET1. No ternary complex is found in cluster B1 for dBET1. **(B)** dBET23. **(C)** dBET57. **(D)** dBET70. Cluster A1 (red) and Cluster B1 (yellow). Black arrows indicate the hinge loops in both DDB1 and Rbx1 proteins. CUL4A (white), NEDD8 (magenta), E2 (ice blue), Ub (light yellow), and Rbx1 (rose).

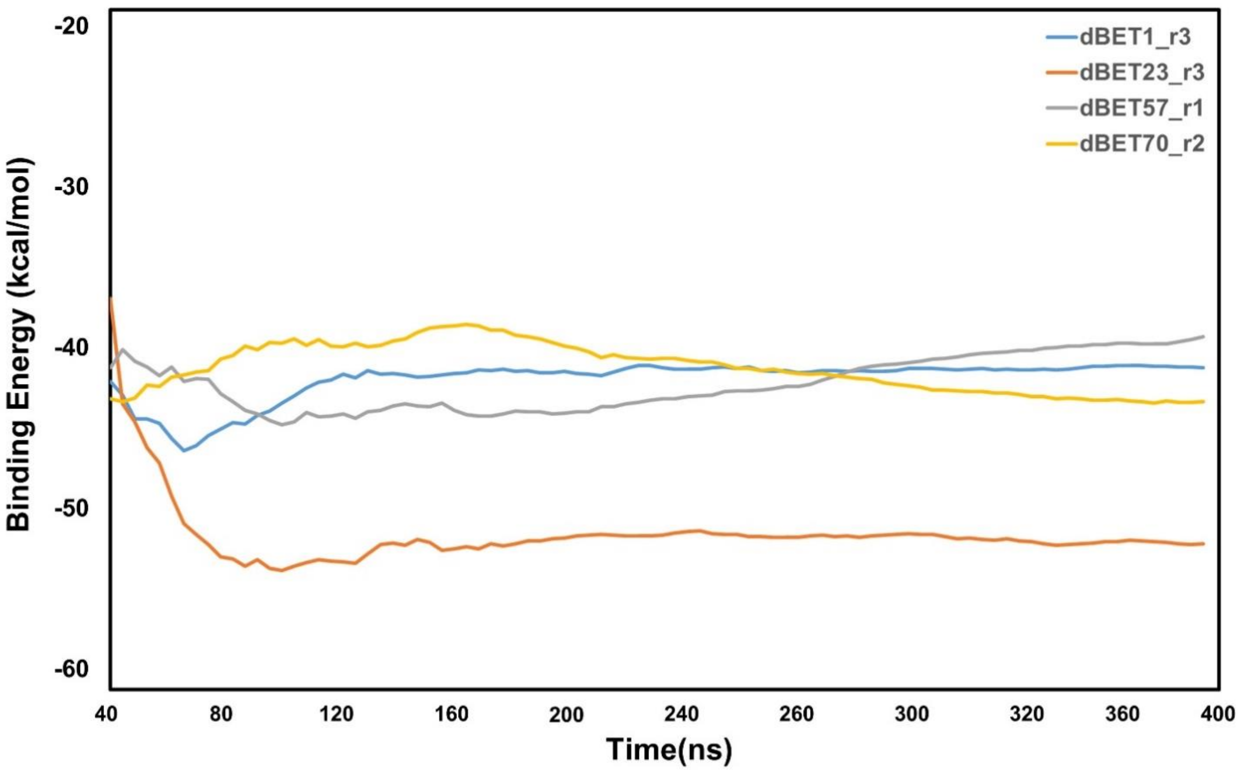


**Figure S10**: **Analysis of protein-protein interaction energies of ternary complexes for each dBETs.** Computed nonbonded interaction energy between proteins (CRBN-BRD4$^{BD1}$) and different dBETs using the MD runs of our selected ternary complexes. (See Method) Only the lowest interaction energy among the 3 random seeds MD simulations is reported here. Note that the energy calculations focus on non-bonded intermolecular interactions and solvation free energy calculations using MM/PBSA, where the configuration entropy loss during protein binding was not explicitly included.

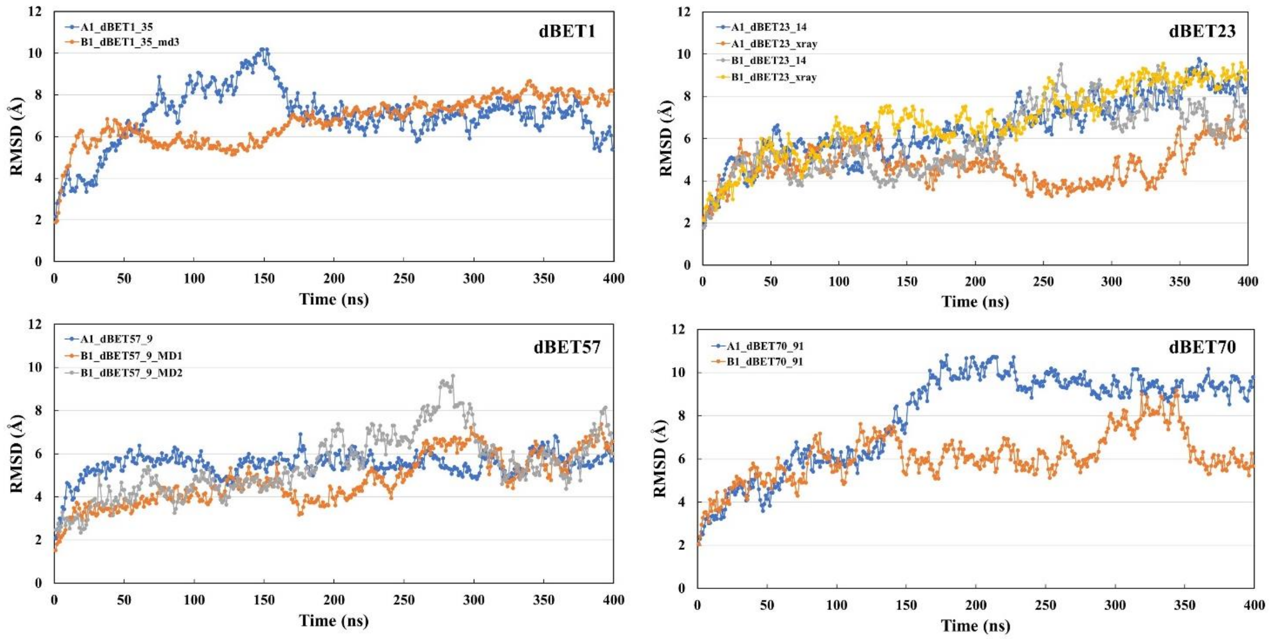


**Figure S11: Root-mean-squared deviation (RMSD) analysis for each degradation machinery complexes.** The RMSD was calculated based on the backbone atoms and the first frame of the trajectory was used as reference.

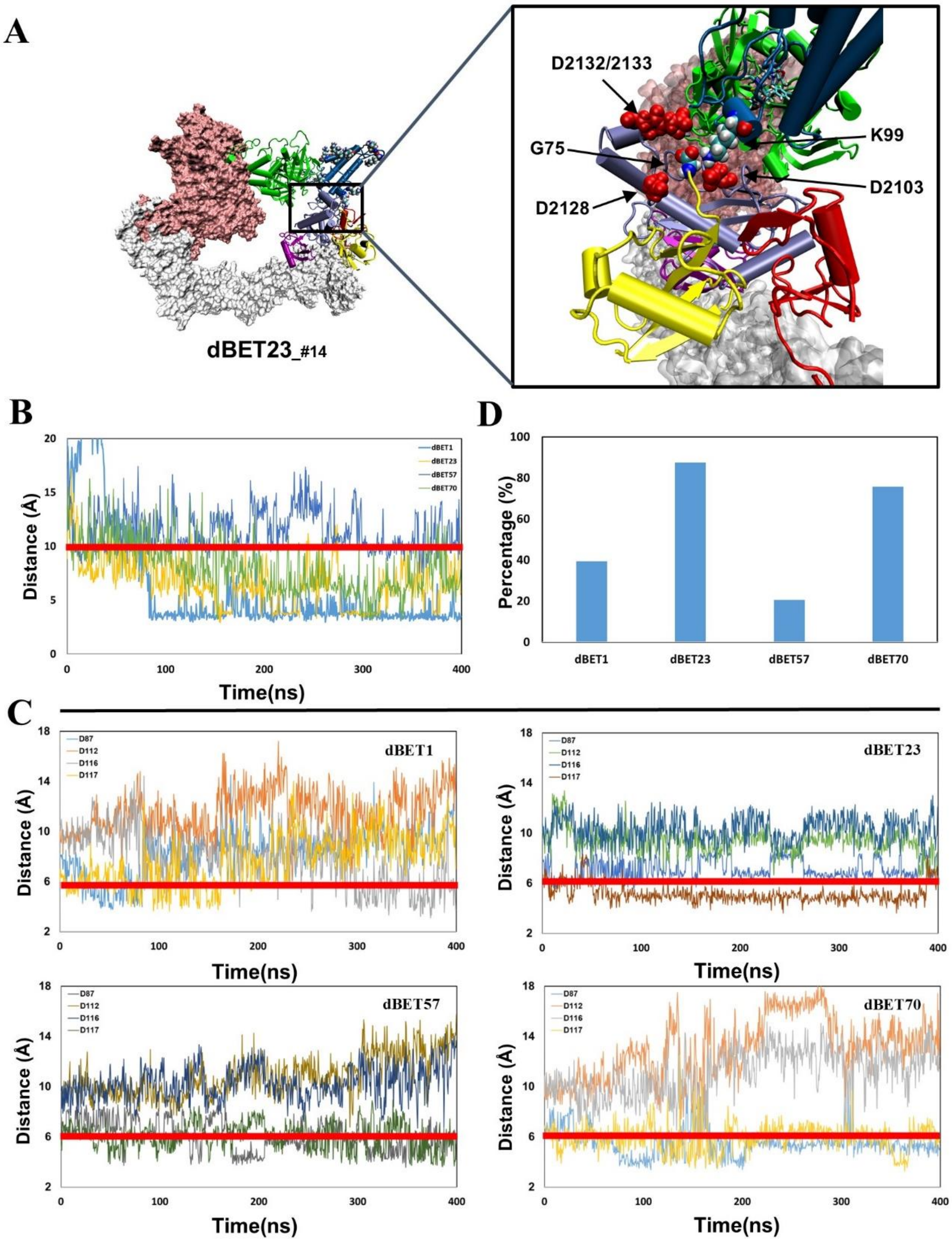


**Figure S12. Quantifying the probability of isoppetide bond formation using two criteria: (1) Lys (N atom) Gly (C atom) distance and (2) Asp (O atom) and Gly (O atom) distance. (A)** Visualizing four Asp of E2 in close proximity of G75 of Ub. Asp is colored in red with vdW representation**. (B)** The Lys (N atom) and Gly (C atom) distance of each dBETs. **(C)** All four Asp (O atom) and Gly (O atom) distance of each dBETs. **(D)** Probability of satisfying both criteria for each dBETs. Red line indicates the cutoff distance for each criterion: 10 Å for Lys (N atom) Gly (C atom) distance; 6 Ã for Asp (O atom) and Gly (O atom) distance.

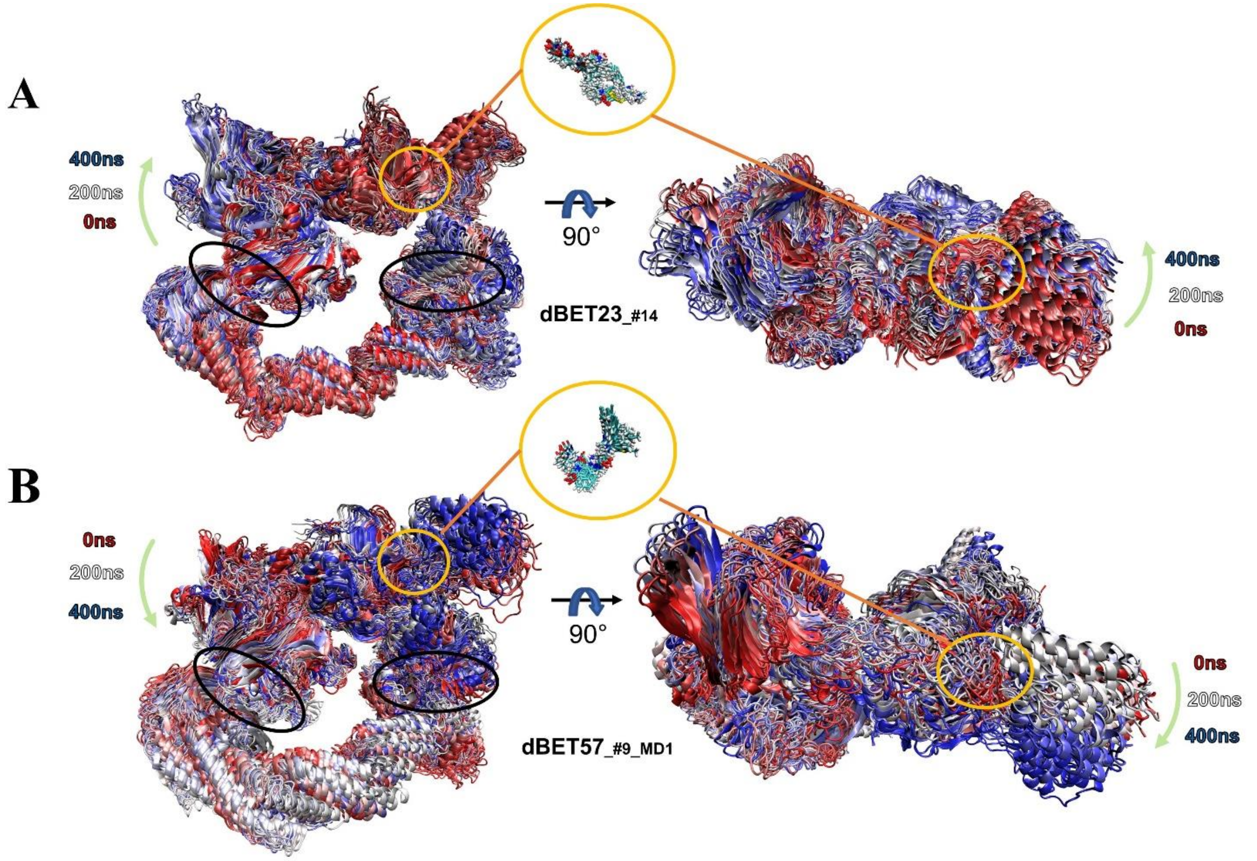


**Figure S13: Dynamic nature of degradation complex.** Overlay of 25 frames from 400-ns MD simulation segment showing the overall dynamic nature of the degradation complex. The green arrows indicate the motion of the hinge region (left panel) and BRD4 (right panel). The whole 400-ns MD trajectories are superimposed based on the backbone of protein. The black circles indicate the two hinge regions: loops in Rbx1 and DDB1 proteins. **(A)** PROTAC dBET23$_{_\#14}$ degradation complex. **(B)** PROTAC dBET57$_{_\#9_MD1}$ degradation complex. Zoom-in representations show the PROTAC dynamics.

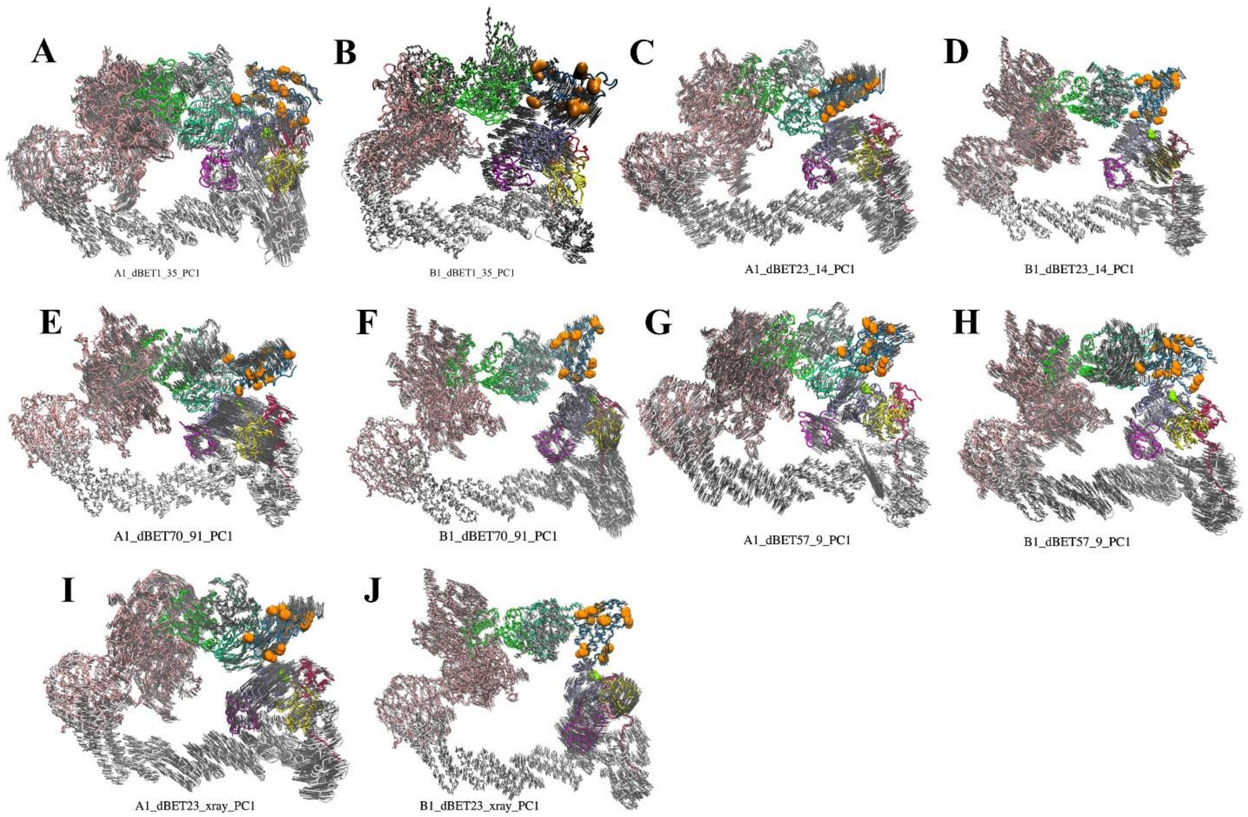


**Figure S14. First PC mode of the predicted ternary complexes. (A-E)** cluster A1 degradation machinery complexes. **(F-J)** cluster B1 degradation machinery complexes. DDB1, pink; CUL4A, white; NEDD8, magenta; E2, ice blue; ubiquitin, yellow; Rbx1, red; HBD of CRBN E3, green; NTD of CRBN E3, mint; CTD of CRBN E3, dark gray; BRD4$^{BD1}$, blue. Orange balls indicate Lys resides on BRD4$^{BD1}$. Light gray arrows indicate the direction of protein movements in the PCA.

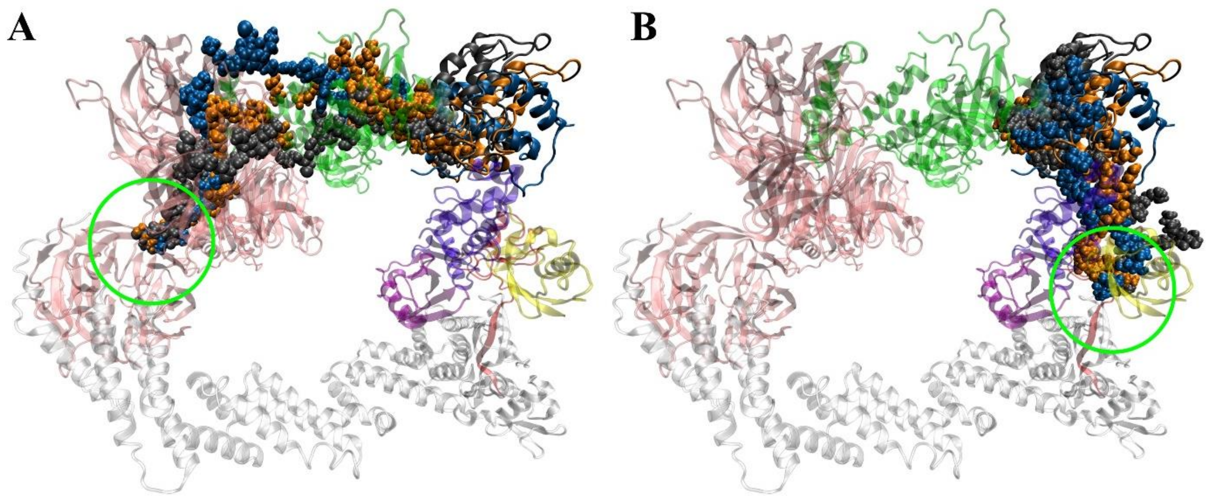


**Figure S15. Correlation between PROTACs and hinge motion.** Tracing the shortest path that propagates through the non-bonded interaction network originated from the linker to the hinge region. (A) Linker to DDB1 (B) Linker to Rbx1. dBET23 (orange), dBET70 (gray), dBET57 (blue), hinge region (green circle).

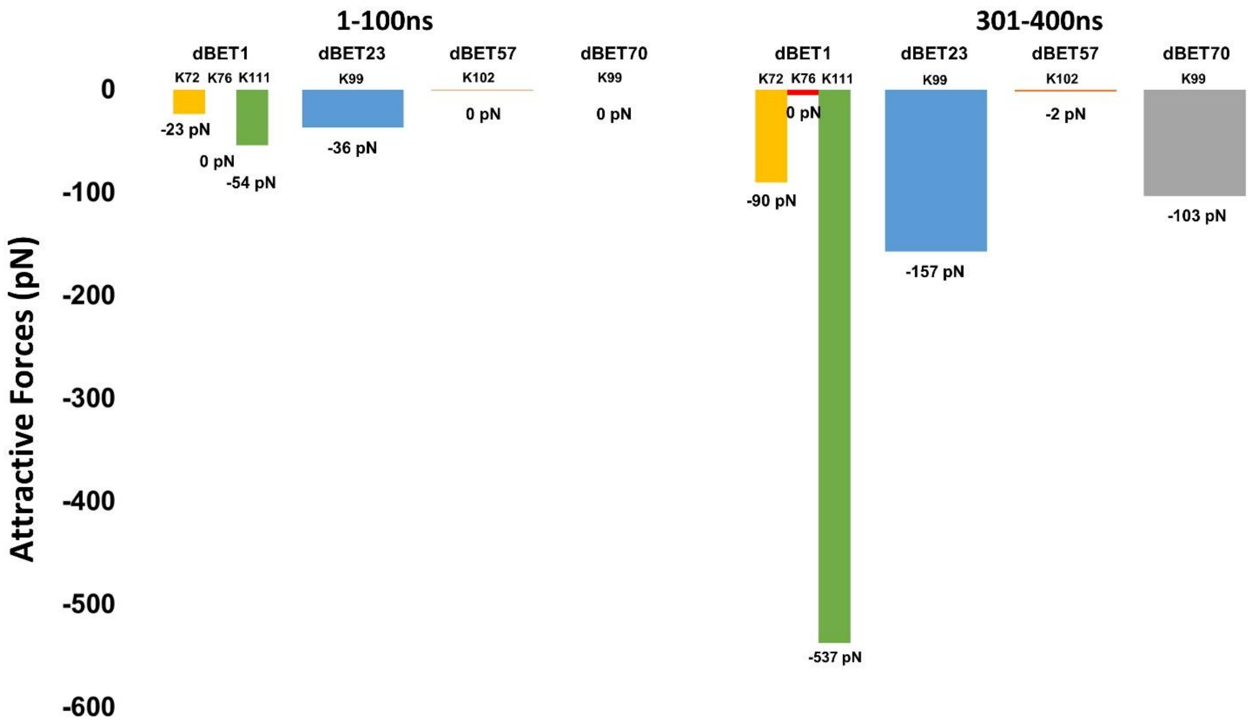


**Figure S16. Attractive force between Lys of BRD4$^{BD1}$ and Gly of Ub.** dBET1 uses Lys72/76/111; dBET23 and dBET70 use Lys99; dBET57 uses Lys102 for Lys-Gly interaction, respectively.

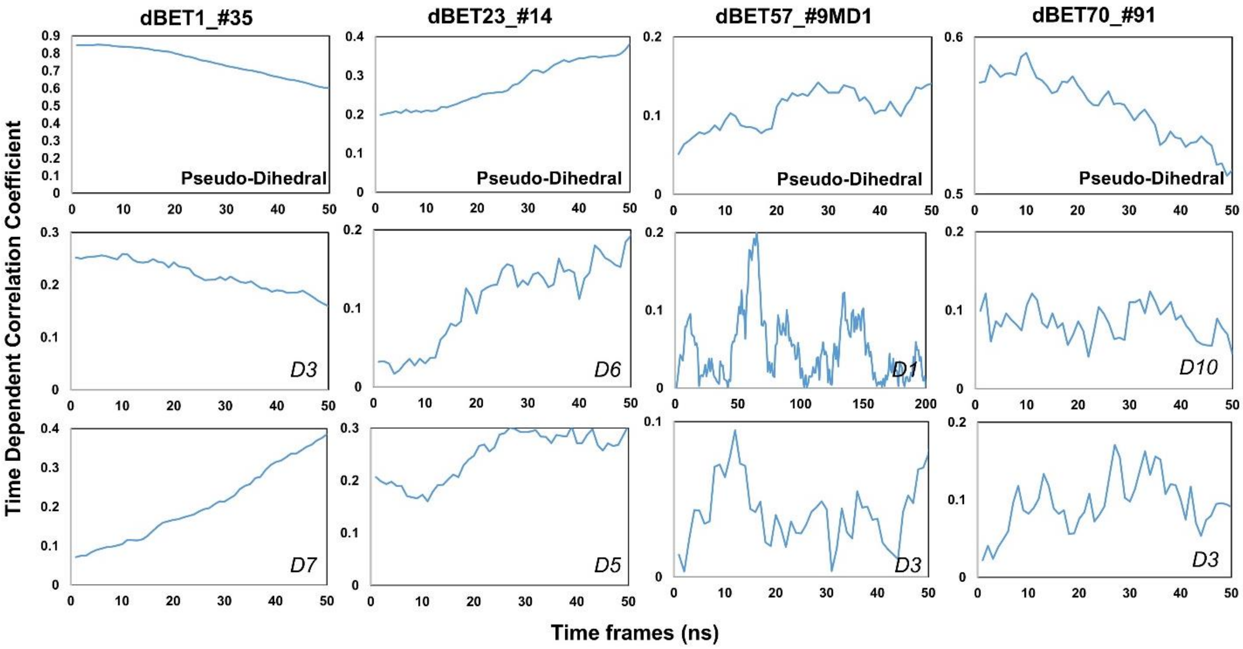


**Figure S17. Time dependent correlation of Psuedo dihedral angles and PROTACs dihedral angles.** (Please refer to Figure S17 for the dihedral index)

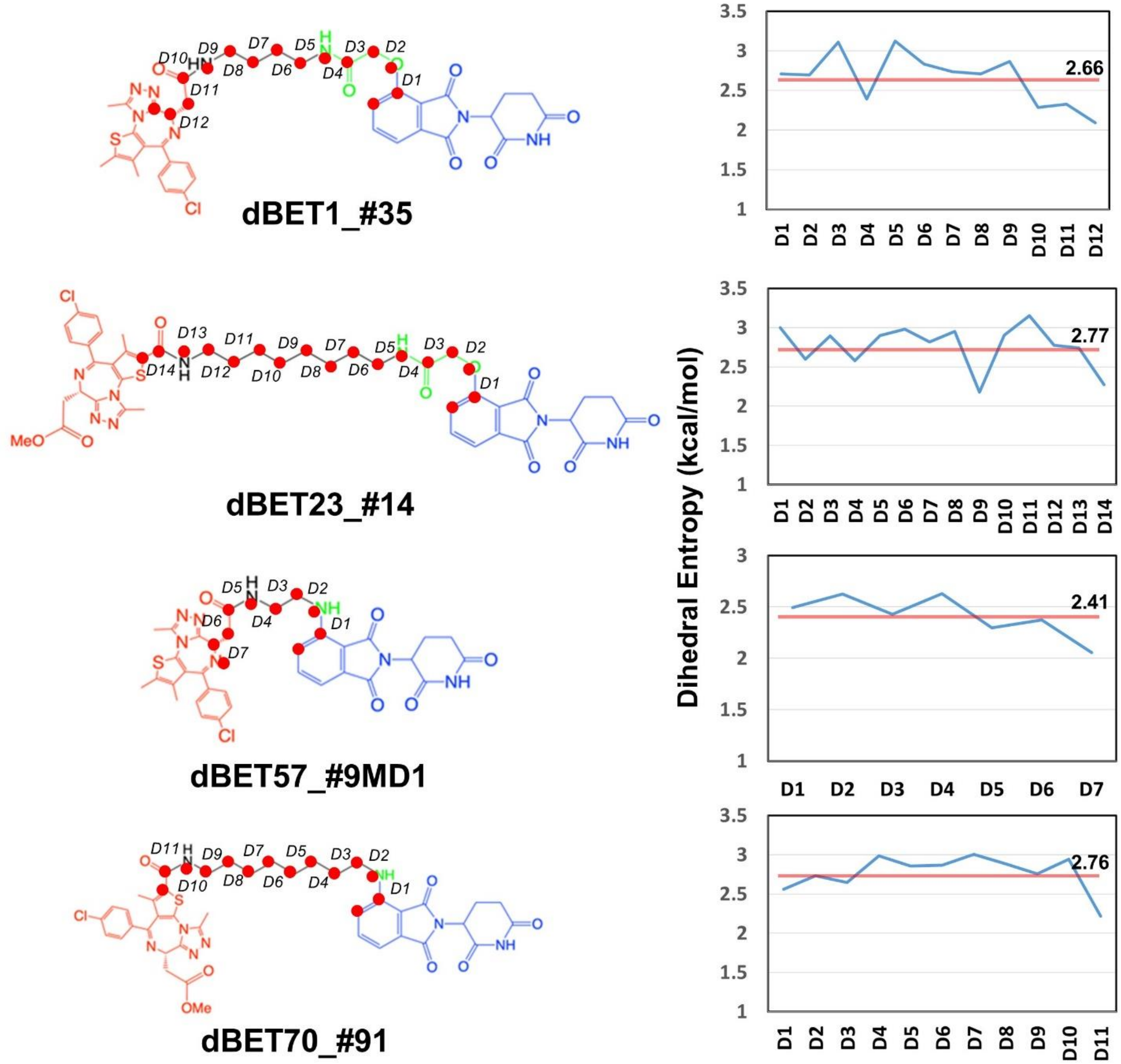


**Figure S18. Dhedral entropies of each dihedra langles of the PROTACs using Gibbs entropy formula.** The probability distribution is the probability distribution of dihedra langles. Larger entropies means the molecules is more flexible and lower values indcates a more rigid molecules.

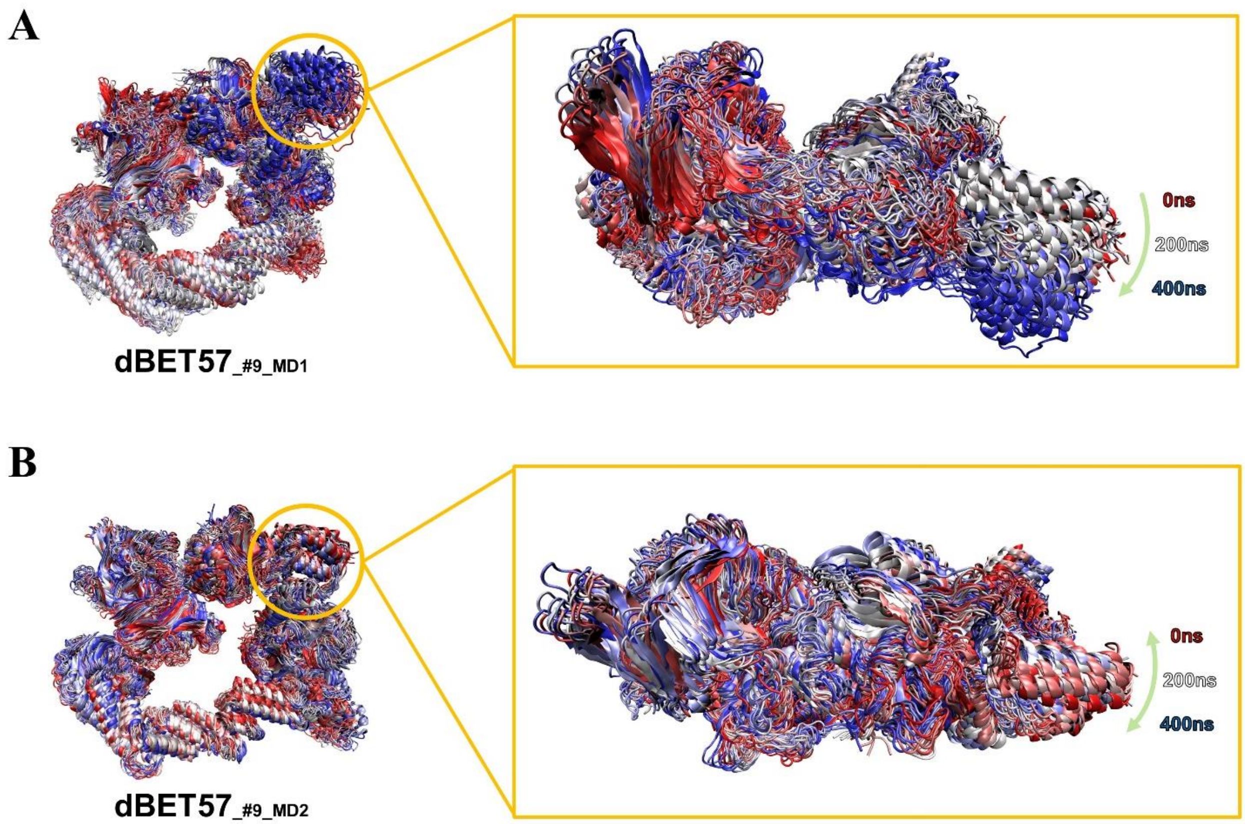


**Figure S19.** Comparing degradation machinery complexes' dynamic of dBET57$_{_\#9_MD1}$ and dBET57$_{_\#9_MD2}$ with different BRD4$^{BD1}$ initial conformations**.** **(A)** BRD4$^{BD1}$ of dBET57$_{_\#9_MD1}$ shows larger motion (10-15 degrees) which eventually leads to ubiquitination. **(B) B**RD4$^{BD1}$ of dBET57$_{_\#9_MD1}$ shows very little motion (~0 degree) and did not recruit Lys for ubiquitination.

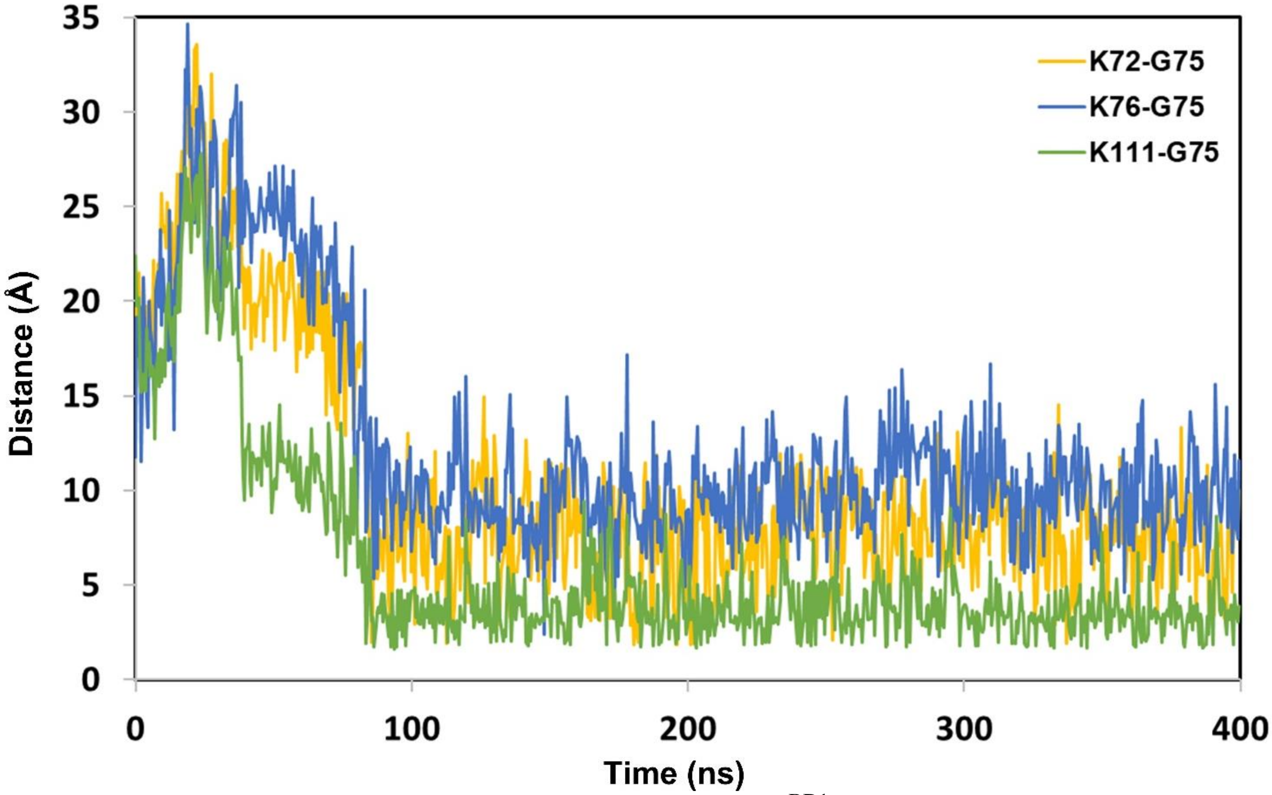

**Figure S20.** The distance between Lys (N atom) of BRD4$^{BD1}$ and Gly (C atom) of Ub from dBET1$_{_\#35}$ degredation complex.

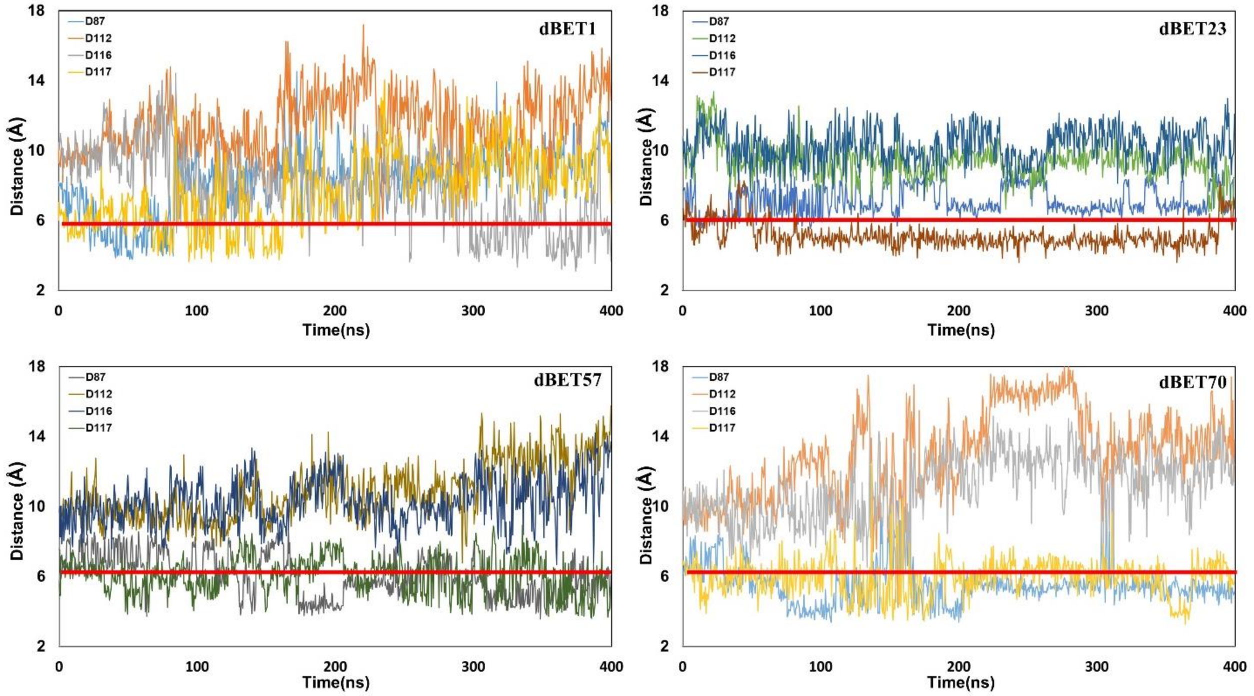


**Figure S21: Lys (N atom) Gly (C atom) distance and (2) Asp (O atom) and Gly (O atom) distance for second seeds of MD simulation.** All four Asp (O atom) and Gly (O atom) distance of each dBETs.

**Table S1:** Summary of the predicted CRBN-dBETx-BRD4$^{BD1}$ ensembles from protein-protein docking. Total ternary conformations were recorded, and conformations with clashes were carefully examined and manually removed.

| PROTAC | $DC_{50/5h}$ (nM)[21] | Degrader Conf. | Total ternary Conf. predicted | Selected reasonable ternary complexes | Reasonable ternary complexes (%) |
|---|---|---|---|---|---|
| dBET1 | ~500 | 125 | 297 | 186 | 62.3 |
| dBET23 | ~50 | 212 | 250 | 168 | 67.2 |
| dBET57 | ~500 | 25 | 40 | 24 | 60.0 |
| dBET70 | 5 | 143 | 217 | 183 | 84.4 |

**Table S2:** Binding scores predicted by MOE protein-protein docking for the CRBN-dBETx-BRD4$^{BD1}$ ternary complexes.

| dBET1 Conf. index # | $E_{total}$ (kcal/mol) | dBET23 Conf. index # | $E_{total}$ (kcal/mol) | dBET57 Conf. index # | $E_{total}$ (kcal/mol) | dBET70 Conf. index # | $E_{total}$ (kcal/mol) |
|---|---|---|---|---|---|---|---|
| **35** | **-7353.1** | 206 | -7272.9 | 22 | -7265.3 | **91** | **-7330.9** |
| 10 | -7352.0 | **14** | **-7221.1** | 24 | -7265.2 | 85 | -7326.2 |
| 27 | -7349.2 | 165 | -7193.7 | 26 | -7264.9 | 90 | -7323.8 |
| 30 | -7345.6 | 168 | -7183.9 | **9** | **-7237.8** | 117 | -7319.3 |
| 101 | -7304.9 | 166 | -7177.8 | 23 | -7225.3 | 88 | -7314.0 |
| 110 | -7279.3 | 217 | -7141.8 | 1 | -7216.9 | 118 | -7308.2 |
| 36 | -7255.5 | 167 | -7141.4 | 7 | -7201.4 | 120 | -7303.2 |
| 106 | -7236.1 | 16 | -7045.1 | 4 | -7153.0 | 89 | -7297.7 |
| 114 | -7260.9 | | | 2 | -7145.9 | 119 | -7295.4 |
| 119 | -7142.3 | | | 3 | -7130.2 | 122 | -7288.8 |
| | | | | 6 | -7125.6 | 125 | -7256.5 |
| | | | | 11 | -7065.6 | 123 | -7254.4 |
| | | | | | | 98 | -7244.7 |
| | | | | | | 121 | -7237.9 |
| | | | | | | 87 | -7219.4 |
| | | | | | | 95 | -7214.7 |
| | | | | | | 96 | -7195.2 |
| | | | | | | 124 | -7104.8 |
| | | | | | | 126 | -7040.4 |

**Table S3:** Accessible lysine residues identified in the CRL4A E3 ligase scaffolds for each PROTAC in each cluster.

| PROTAC | A1 | A2 | A3 | A4 | A5 |
|---|---|---|---|---|---|
| dBET1 | 55,57,102,112 | 55,57,112 | 112,155 | 112 | 55,57,72,91,111, 112,155,160 |
| dBET23 | 55,102,141,15 5,160 | 55,99,102,112,141, 155,160 | 72,76,111,155 | N/A | 155,160 |
| dBET57 | 55,112 | 55,112 | N/A | 111,112 | 111,112 |
| dBET70 | 55,141,160 | 55,99,102,112,141, 155,160 | 76,155,160 | N/A | 155,160 |

| PROTAC | B1 | B2 | B3 | B4 |
|---|---|---|---|---|
| dBET1 | 57,55,72,102,111,112 | 55,57,72,102,111,112,160 | N/A | N/A |
| dBET23 | 55,57,99,102,112 | 55,57,99,102,112 | N/A | N/A |
| dBET57 | 57,55,111,112 | 55,57,72,111,112 | N/A | N/A |
| dBET70 | 55,57,99,102,112 | 55,57,99,102 | N/A | N/A |

**Table S4:** PCA analysis with the first two PCs projection coverage in each degradation machinery complex. The essential motions in the first two PCs of ten degradation machinery complexes were calculated based on the backbone atoms. The coverage of the first two PCs were reported. Note, all, expect A1_dBET1 $_{\#35}$ and A1_dBET57 $_{\#9}$ have the first two PCs coverage is below 60%.

| | A1 | | | B1 | | |
|---|---|---|---|---|---|---|
| PROTAC | PC1 | PC2 | PC1+PC2 | PC1 | PC2 | PC1+PC2 |
| dBET1 $_{\_\#35}$ | 49.2% | 9.0% | 59.2% | 69.0% | 9.6% | 78.6% |
| dBET23 $_{\_\#14}$ | 47.3% | 13.4% | 60.7% | 64.8% | 7.2% | 72.0% |
| dBET57 $_{\_\#9}$ | 28.4% | 13.8% | 42.2% | 49.8% | 10.8% | 60.6% |
| dBET70 $_{\_\#91}$ | 66.2% | 8.6% | 74.8% | 51.1% | 12.3% | 63.4% |
| dBET23 $_{\_xray}$ | 50.7 % | 11.9% | 62.6% | 50.7% | 15.4% | 66.1% |
